# Cold-acclimation, not motor inactivity, attenuates GABA signaling in the respiratory network of bullfrogs in response to overwintering

**DOI:** 10.64898/2026.08.17.745239

**Authors:** Renato Filogonio, Hafsa Yaseen, Joseph M. Santin

## Abstract

Neural circuits produce reliable activity even after environmental disturbances. This occurs because neurons respond to perturbations in a compensatory manner, a process termed homeostatic plasticity. Bullfrogs undergo prolonged periods underwater during winter, when lung ventilation and its neural control system ceases activity, but air-breathing resumes unscathed when environmental temperatures increase weeks to months later. Compensatory neural mechanisms that contribute involve upregulation of excitatory synaptic transmission on motoneurons driven by inactivity, but whether inactivity or acclimation to low temperatures drive other forms of compensation is not known. The GABA_A_ receptor contribution to respiratory rhythm generation is downregulated following overwintering, which promotes network excitability. Therefore, we disentangled the contributions of cold temperature acclimation and inactivity experienced during overwintering on reduced GABAergic signaling. Here, we show that cold temperature, and not inactivity, reduces GABA_A_ signaling in the respiratory rhythm generating network, without influencing GABA_A_ transmission onto motoneurons. Therefore, cold temperature acclimation drives reduced GABAergic signaling selectively in inter-neuronal rhythm generating circuits, while excitatory motoneurons synapses are strengthened by inactivity in the overwintering environment. Most work interprets compensatory plasticity as activity-dependent during activity perturbations, but we reveal that different aspects of a disruptive environment elicit distinct forms of plasticity across a motor network.

## Introduction

Changing environmental conditions (*e*.*g*. temperature, acidity, salinity, oxygen availability) tend to destabilize the organism’s internal environment and therefore, its function. It is widely recognized that organisms use various sets of feedback mechanisms throughout the body that compensate for internal variation to maintain physiological function (*i*.*e*. homeostasis). Neural circuits are thought to sense changes and regulate activity during perturbations by adjustments in synaptic strength and intrinsic membrane properties. This elicits compensatory changes in excitation or inhibition that aim to maintain relatively stable neural physiological function, termed ‘homeostatic plasticity’ [1,2]. While most studies indicate that homeostatic regulation during activity perturbations are dependent on activity sensing and transduction [1,3–5], concomitant abiotic environmental conditions may also affect network activity. For example, during winter cold temperatures may induce hibernation where body temperature and energy requirements are reduced for prolonged periods [6]. Consequently, low temperatures reduce metabolism, and therefore the demands for many neuronal functions, which could each in principle act as signals for plasticity. Therefore, the degree to which reduced neural activity vs. factors associated with a complex environment that ultimately reduce neural activity in the first place serve as signal for compensation can be obscure.

An extreme example of this scenario occurs in the respiratory network of American bullfrogs and other ranid frogs that inhabit northern latitudes. Lung ventilation is necessary for metabolic homeostasis; however, at cold temperatures, gas exchange requirements can be fully provided by cutaneous respiration, allowing frogs to stay underwater for long periods without requiring air-breathing [7]. Therefore, during hibernation, the respiratory circuits responsible for producing lung ventilation are silent [8]. Consistent with homeostatic theory [9,10], inactivity serves as a key signal that enhances excitability during hibernation by upregulating excitatory AMPA-glutamate receptors on motoneurons [11–13]. Interestingly, the enhancement of AMPA receptors on motoneurons correlated with greater evoked synaptic transmission selectively at cool temperatures, indicating that homeostatic regulation can have broader effects on network robustness than globally enhancing excitability [13]. Therefore, in a complex environment with multiple interacting variables, inactivity seems to elicit homeostatic adjustments in neural properties that improve performance across a range of environmental conditions.

While homeostatic tuning rules upregulate excitatory synaptic receptors, it is not clear if these same processes also control other forms of compensation that arise during hibernation. For example, inhibitory signaling in neural circuits are mainly orchestrated by ligand-gated Cl^-^ channels activated by the neurotransmitter γ-aminobutyric acid (GABA) through GABA_A_ receptors [14]. After overwintering, the respiratory network has a reduced contribution of GABA_A_ receptors to rhythm generation. In this way, the application of GABA_A_ receptor antagonist typically stops respiratory-related activity in controls, while it had no effect after hibernation. The loss of GABA_A_ signaling in hibernators had the unexpected consequence of enhancing excitability at cool temperatures by removing inhibition that is stimulated during cooling to slow activity in controls [15], a surprising parallel to enhanced transmission at excitatory motoneuron synapses due to upregulated receptors through homeostatic mechanisms [13]. These results suggest that both AMPA receptors and the GABA system may be modulated by inactivity during hibernation. However, it remains possible that GABA may be controlled independently. In cultured cortical neurons, excitatory synapses and intrinsic excitability were enhanced during inactivity by mechanisms involving calcium/calmodulin kinase, while GABA inhibition was decreased via separate and undetermined mechanisms [16]. In addition, circulating levels of GABA in warm-acclimated epaulette sharks (*Hemiscyllium ocellatum*) are comparatively higher than cold-acclimated sharks [17], indicating that temperature *per se* may play a role in the regulation of the GABA system. Therefore, we investigated what stimulus (i.e. cold-acclimation or inactivity during overwintering conditions) reduces GABA_A_ signaling in the central pattern generator of the bullfrogs. We also examined whether those stimuli affect GABA_A_ signaling on motoneurons using whole cell patch-clamp to measure evoked inhibitory postsynaptic currents (IPSC). This allowed us to discriminate the location of GABAergic compensatory plasticity within an ecologically meaningful context of the overwintering environment.

## Material and Methods

### Animal acquisition and maintenance

Adult American bullfrogs (*Aquarana catesbeiana*) were acquired from Niles Biological (Sacramento, CA, USA) and maintained in the facilities from the Division of Biological Sciences of the University of Missouri (Columbia, MO). Frogs were randomly assigned to one of three experimental groups which were kept in plastic tanks with dechlorinated water continuously bubbled with room air, and a 12:12h light:dark cycle. The control group was kept at room temperature (∼22°C) and fed once a week with pellets provided by the seller. Two groups of cold-acclimated frogs (ventilating and overwintered) were kept at a temperature-controlled room set to 20°C for approximately 3 days whereafter room temperatures were reduced by 2°C per day until it reached 4°C. One group of cold-acclimated frogs were kept with a layer of water covering half of their body, thus allowing lung ventilation, whereas for the overwintering group plastic nets were placed in the water level as soon as room temperature reached 4°C, which impeded lung ventilation. Buccal pumping in the ventilating group was visually confirmed upon daily animal welfare checks. Cold-acclimated frogs from both ventilating and overwintered groups were maintained at least 2 weeks under 4°C before commencement of experiments, which was sufficient to increase the amplitude of spontaneous excitatory postsynaptic currents in this species [12]. All procedures were approved by the Animal Care and Use Committee from the University of Missouri (protocols #39264 and #65623).

### Tissue preparation

Frogs were initially anesthetized with a cotton soaked in isoflurane (1 ml) within a plastic box of approximately 1l. After loss of pedal reflexes, frogs were decapitated and the head was transferred to a cold artificial cerebrospinal fluid solution (aCSF, in mmol^-1^: 104 NaCl, 4 KCl, 1.4 MgCl_2_, 7.5 D-glucose, 40 NaHCO_3_, 2.5 CaCl_2_, 1 NaH_2_PO_4_, gassed with 1.5% CO_2_/98% O_2_ for pH = 7.85). The brainstem-spinal cord was dissected free and had the dura removed.

### Extracellular protocol

For the extracellular recordings, the brain-spinal cord with the nerves intact was moved to a dish perfused with aCSF at a flow rate of 6 ml × min^-1^ with a peristaltic pump (Ranin Rabbit 4 channel head, Mettler Toledo Rainin, Oakland, CA, USA). Suction electrodes were used to measure the respiratory burst activity from the hypoglossal rootlet. We measured nerve activity in this condition during 1 h before applying the GABA_A_ receptor antagonist bicuculline (10 μM) for 30 min (Fig. 1).

**Figure 1.**
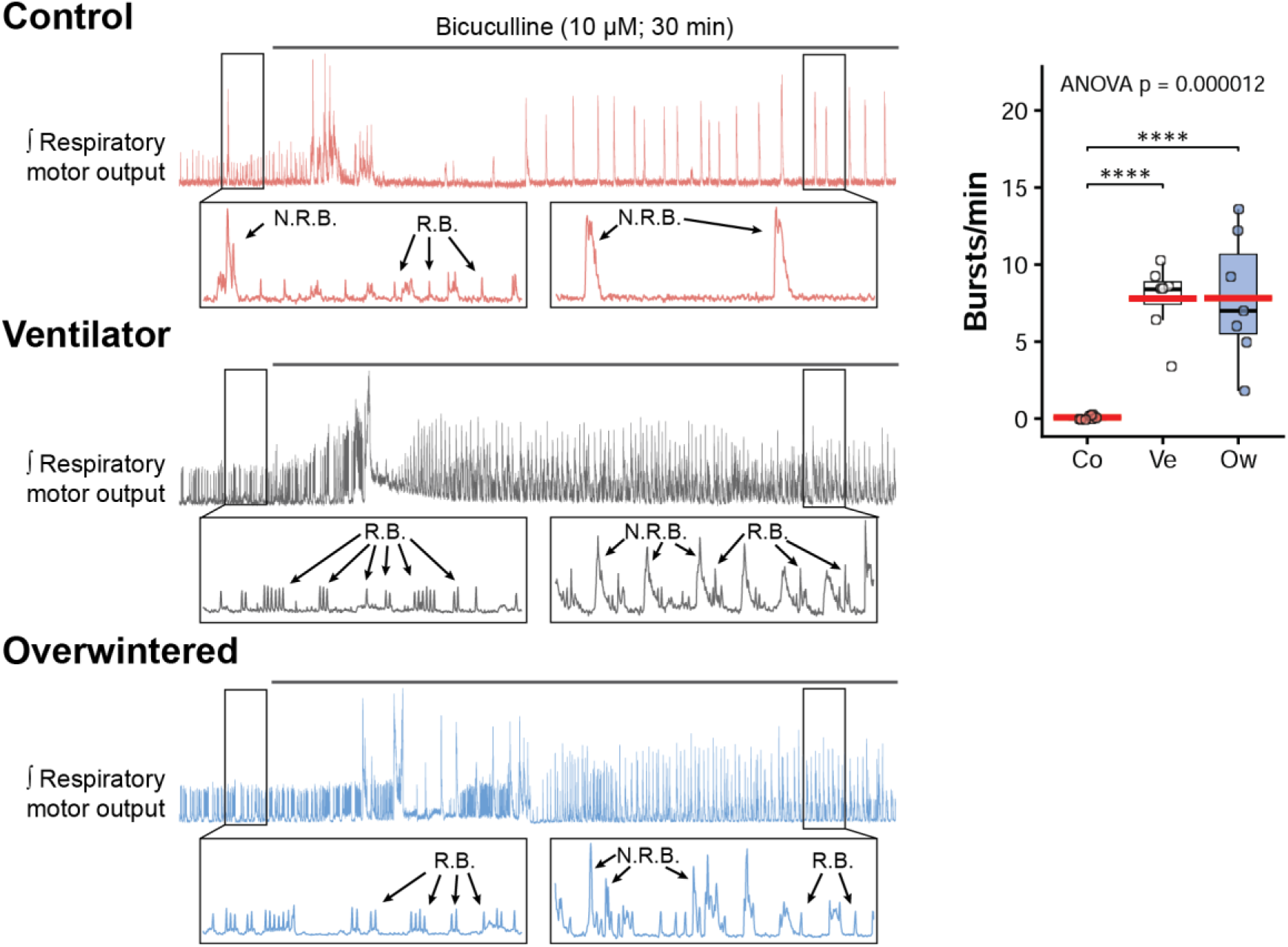
Effects of bicuculline on the brainstem respiratory network activity of the American bullfrog (*Aquarana catesbeiana*). On the left, original recordings of the brainstem motor activity of control (magenta), ventilator (gray) and overwintered (blue) groups before and after 30 min of exposure to the GABA_A_ antagonist, bicuculline (10 μM). The recordings show both respiratory bursts (R.B.) and non-respiratory bursts (N.B.R.). On the right, the boxplot graph compares the brainstem respiratory motor activity (bursts/min) after 30 min of exposure to bicuculline in the control (Co; magenta circles; n = 8), ventilator (Ve; white circles; n = 7), and overwintered (Ow, blue circles; n = 7) groups. The horizontal red lines over the boxplots indicate the mean values. Groups were compared with a one-way ANOVA and pairwise comparisons performed with a Holm-Šydák *post hoc* test. Significance level was p < 0.05 ‘*’; p < 0.01 ‘**’; p < 0.001 ‘***’; p < 0.0001 ‘****’.

### Electrophysiological protocol

For the electrophysiology experiments, after dissection the brain-spinal cord was glued to an agar block and sliced (300 μm thick) with a vibrating microtome (Technical Products International Series 1000, St. Louis, MO, USA). Voltage clamp experiments were conducted using the same equipment previously described [12]. Thin walled glass pipettes (resistance: 3-8 MΩ) were fabricated using a Sutter Instruments puller (model P87, Novato, CA, USA) and filled with an intracellular solution (in mM: 95 CsCl, 2 MgCl_2_, 10 HEPES, 1 Na_2_ATP, 0.1 Na_2_GTP, 10 EGTA, 1 CaCl_2_, 10 TEA-Cl; pH = 7.2 corrected with CsOH; [18] prepared with Milli-Q water. Slices were continuously bathed with a solution of aCSF containing the glycine receptor antagonist, strychnine (5 μM), and the ionotropic glutamate receptor antagonist, DNQX (20 μM), bubbled with 1.5% CO_2_/98% O_2_. To ensure this solution isolated GABA_A_ currents, we validated this protocol by adding the GABA_A_ receptor antagonist, bicuculline (50 μM), to ensure all evoked currents were silenced (Fig. 2A).

**Figure 2.**
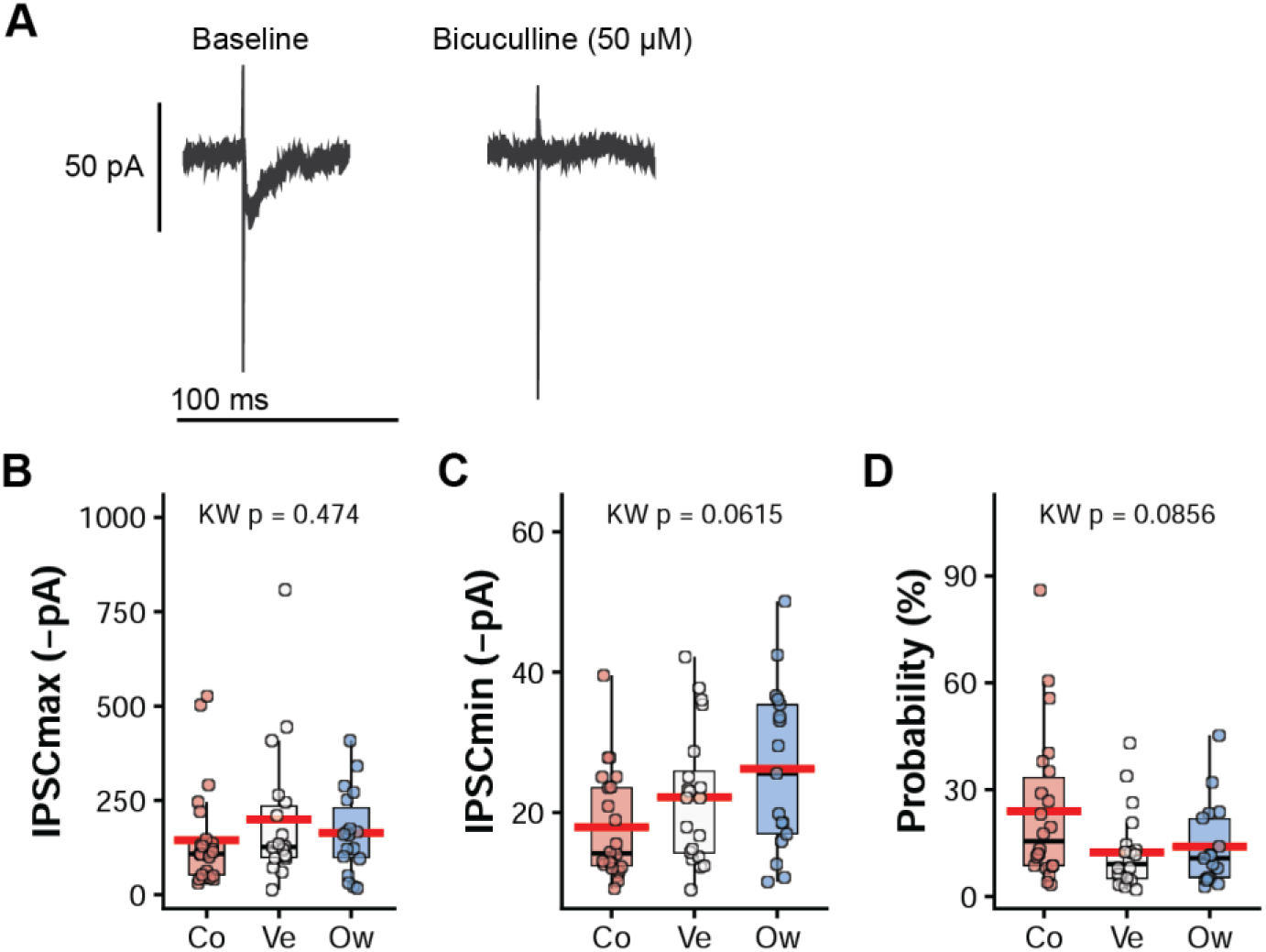
Measurements of the inhibitory postsynaptic currents (IPSC) of the hypoglossal motoneurons from American bullfrogs, *Aquarana catesbeiana*. **A**) Original recordings demonstrating the validation of the protocol. Baseline currents were recorded when cells were in voltage clamp while the slice was bathed with the glutamate antagonist, DNQX (20 μM) and the glycine antagonist, strychnine (5 μM). Complete inhibition was attained by adding the GABA_A_ antagonist, bicuculline (50 M). **B**) Amplitude (-pA) of maximum IPSC (Co: n = 24 from 8 frogs; Ve: n = 20 from 7 frogs; Ow: n = 18 from 6 frogs); **C**) Amplitude (-pA) of minimum IPSC (Co: n = 22 from 8 frogs; Ve: n = 20 from 7 frogs; Ow: n = 17 from 6 frogs); **D**) Probability (%) of evoking an IPSC at minimum stimulation (Co: n = 22 from 8 frogs; Ve: n = 20 from 7 frogs; Ow: n = 17 from 6 frogs). Groups tested were control (Co; magenta circles), ventilating (Ve; white circles), and overwintered (Ow, blue circles) frogs. The horizontal red lines over the boxplots indicate the mean values. Groups were compared with a Kruskal-Wallis test; significance level was p < 0.05.

Inhibitory postsynaptic currents were evoked using a bipolar tungsten electrode (WE3ST0.1B10, Microprobes for Life Sciences, Gaithersburg, MD, USA) placed at the axons of the solitary tract [19]. The electrodes were connected to an isolated pulse stimulator (Model 2100, A–M Systems, Carlsborg, WA, USA). The temperature of the solution bathing the slices was controlled with a bipolar in-line temperature controller (Model CL-100, Warner Instruments, Hamden, CT, USA) and monitored with a thermocouple placed close to the tissue.

Hypoglossal motoneurons, which innervate the buccal floor compressor muscles associated with breathing in amphibians [20] were accessed using whole-cell patch clamp. After the cell was accessed, we determined the maximum inhibitory postsynaptic currents (IPSCmax) by increasing the stimulus currents until we observed the asynchronous vesicle release, after which we reduced the stimulus intensity until we could maintain the largest amplitude without the asynchronous vesicle release [21]. IPSCmax were recorded for 30 s at a frequency of 0.2 Hz. After that, minimum inhibitory postsynaptic currents (IPSCmin) were determined by decreasing the stimulation until no currents were observed, after which stimulation intensity was increased stepwise until the minimum value where we could observe at least one current within 10 consecutive stimulation events [13]. IPSCmin were recorded for 5 min.

### Drugs

Strychnine hydrochloride was purchased from Sigma-Aldrich (St Louis, MO, USA). (-)-Bicuculline methiodide and DNQX disodium salt were acquired from Hello Bio (Princeton, NJ, USA).

### Data analysis and statistics

Bursting frequency (bursts/min) from nerve rootlets was assessed by counting the amount of respiratory bursts during 5 min. We counted respiratory bursts just before applying bicuculline to the bath, and in the last 5 min of an exposure of 30 min long (Fig. 1). IPSCmax (-pA) were the averaged amplitude of three maximum evoked currents. IPSCmin (-pA) were the average of the amplitude of the evoked minimum currents identified with the ‘Peak Analysis’ module from LabChart (v.8.0, ADInstruments). Release probability (%) measures the probability of release of neurotransmitters and was calculated as the percentage of events that evoked an IPSCmin during the five minutes of stimulation.

Differences between groups were tested with a one-way ANOVA followed by a Holm-Šydák *post hoc* test. When data did not comprise with the assumption of normality, differences were tested with a Kruskal-Wallis followed by a Dunn test for pairwise comparisons. All statistics were performed with R v.4.6.0 [22–24]. Data are mean ± SD.

## Results

To disentangle the effects of cold-acclimation and inactivity over the attenuation of GABA_A_ signaling during hibernation in American bullfrogs [15], the experimental groups tested were control (warm-acclimated and respiratory apparatus active), cold ventilators (cold-acclimated and respiratory apparatus active) and overwintered (cold-acclimated and inactive respiratory apparatus). Brainstem bursting rates before bicuculline exposure was similar between groups (control: 6.78 ± 4.51 bursts/min; cold ventilators: 6.61 ± 9.44 bursts/min; overwintered: 7.34 ± 6.94 bursts/min). Exposure of the brainstem to bicuculline (10 μM) for 30 min was sufficient to silence respiratory bursts in the control group (Fig. 1) as previously demonstrated [15]. While there was a cessation of respiratory-related bursting, large amplitude non-respiratory bursts remained [25]. Corroborating previous results [15,21], the respiratory network remained active after the bicuculline exposure amid the non-respiratory bursts in the overwintered group, but this was also observed in the cold ventilators, with no statistical differences between these two groups (Fig. 1).

We also investigated whether the downregulation of GABAergic signaling observed in rhythmic respiratory-related activity was accompanied by alterations of inhibitory GABA_A_ receptor synapses in the hypoglossal motoneurons. The GABA_A_-selective isolation of these evoked currents was confirmed by bicuculline application, which abolished all evoked activity (Fig. 2A). Maximum and minimum IPSC were evoked by stimulating the axons at the solitary tract, and release probability was assessed as the amount of IPSCmin over the stimulation events for 5 min. However, none of those variables were significantly different between treatments (Fig. 2B-D), indicating reductions in GABA_A_ signaling was restricted to the premotor respiratory rhythm generating circuits rather than motoneurons.

## Discussion

Hibernation reduces GABA_A_ signaling in the bullfrog respiratory network [15]. The inhibition of respiratory bursts by bicuculline in *A. catesbeiana* under control conditions, and the maintenance of respiratory bursts in overwintered frogs was previously described [15,21]. Contrary to our expectations based on excitatory motoneuron synapses [13], this reduction in GABA_A_ signaling was localized to the rhythm generating circuits and entirely driven by cold-acclimation, rather than inactivity of the motor system that occurs during hibernation. While both forms of plasticity enhance function of the network at cool temperatures [13,15], these results demonstrate two apparently ‘homeostatic’ network modifications (i.e., those that oppose an activity perturbation) may be controlled by entirely distinct aspects of a complex environment.

The present results support that cold-acclimation, but not inactivity, reduces the reliance on the GABA_A_ system in the bullfrog respiratory network. We make this conclusion because bicuculline exposure silences the respiratory network activity in control (warm and active) frogs whilst both ventilators (cold-acclimated and active) and overwintered (cold-acclimated and inactive) frogs remained operational. Previous studies connect these two processes in other hibernating species. In the ground squirrel (*Citellus citellus*), GABA levels increased in the pons and spinal cord when temperature was experimentally shifted from 5°C to 35°C, initiating the process of arousal from the hibernating state within 1.5-2.5 h [26]. At high concentrations, GABA may desensitize GABA_A_ receptors, ultimately reducing membrane conductance to Cl^-^ [14,27–29]. During our experiments, bullfrogs acclimated to 4°C were dissected and later tested at room temperature of approximately 22°C, which may have elevated the GABA concentration and consequent desensitization of GABA_A_ receptors. On the other hand, the brainstem of 13-lined ground squirrel (*Ictidomys tridecemlineatus*) undergoes seasonal changes in the expression of GABA_A_ receptor subunit composition, which suggests GABA_A_ receptor expression may vary according to environmental temperatures [30]. While the mechanistic relationship between cold-acclimation and the GABA system (e.g., whether receptors are downregulated or presynaptic GABA synthesis is reduced) remains to be explored, our results nevertheless are consistent with the idea that inactivity does not drive these responses.

Synaptic inhibition of premotor neurons are governed by GABA_A_ and glycine receptors coupled with Cl^-^ ionotropic channels, and are pivotal for function of rhythmic oscillatory networks [31–35]. Contrary to mammals, where blockade of GABA_A_ receptors does not stop respiratory networks [36], the current data, in accordance with previous observations [15,21,37] unequivocally demonstrate that GABA_A_ receptors are crucial in generating ventilatory rhythm for lung ventilation in adult amphibians. Blockade of glycine receptors also silences respiratory bursts in *A. catesbeiana* [37]. Since respiratory network activity persists during the application of bicuculline in both ventilators and overwintered bullfrogs in the present study, it is possible that cold-acclimation somehow causes glycine receptors to play a more dominant role in generating the respiratory rhythm. Alternatively, pacemaker neurons dependent on persistent Na^+^ currents or Ca^2+^-activated nonspecific cationic currents were hypothesized as a mechanism accounting for rhythmogenesis [32,33,38,39]. In *A. catesbeiana*, it was suggested that lung-related bursts of tadpoles switch their rhythmogenic mechanism from pacemaking to network inhibition when they metamorphose to adults [40]. Likewise, the perspective of bullfrogs switching between different mechanisms of rhythm generation in response to cold-acclimation is intriguing and merit further attention.

## Conclusions

The present results demonstrate that during hibernation, cold-acclimation is responsible for the changes regarding inhibitory GABAergic signaling in the respiratory network of the American bullfrog, *A. catesbeiana*. Additionally, inhibitory synapses in the hypoglossal motoneuron are unaffected during overwintering, which is in stark contrast to AMPA-glutamate excitatory synapses where inactivity during overwintering is the main factor leading to synaptic plasticity [13]. Therefore, excitatory and inhibitory synapses in different parts of the network respond to different components of the same environmental stressors during hibernation. This is in accordance with previous observations suggesting independent regulatory mechanisms between excitatory and inhibitory synaptic scaling [16]. These results highlight that the plethora of environmental inputs received by an organism elicit various and complex mechanisms driving homeostatic responses in an ecologically realistic context.

## Acknowledgements

The authors thank Natalie Heath and Nikolaus Bueschke for help with animal caretaking.

## Funding

J.M.S. was funded by the National Institutes of Health (R01NS114514).

## Notes

https://doi.org/10.6084/m9.figshare.32942075

### Competing Interest Statement

The authors have declared no competing interest.

